# MeCP2 regulates Gdf11, a dosage-sensitive gene critical for neurological function

**DOI:** 10.1101/2022.10.05.510925

**Authors:** Sameer S. Bajikar, Ashley G. Anderson, Jian Zhou, Mark A. Durham, Alexander J. Trostle, Ying-Wooi Wan, Zhandong Liu, Huda Y. Zoghbi

**Author notes:** Corresponding author: Huda Y. Zoghbi; address: Jan and Dan Duncan Neurological Research Institute, 1250 Moursund Street, Suite 1350, Houston, TX, 77030.

## Abstract

Loss- and gain-of-function of MeCP2 causes Rett syndrome (RTT) and *MECP2* duplication syndrome (MDS), respectively. MeCP2 binds methyl-cytosines to finely tune gene expression in the brain, though identifying genes robustly regulated by MeCP2 has been difficult. By integrating multiple transcriptomics datasets, we identified that MeCP2 finely regulates Growth differentiation factor 11 (*Gdf11). Gdf11* is down-regulated in RTT mouse models and is inversely up-regulated in MDS mouse models. Strikingly, genetically normalizing *Gdf11* dosage levels improved several behavioral deficits in a mouse model of MDS. Next, we discovered that losing one copy of *Gdf11* alone was sufficient to cause multiple neurobehavioral deficits in mice, most notably hyperactivity and decreased learning and memory. This decrease in learning and memory was not due to changes in proliferation or numbers of progenitor cells in the hippocampus. Lastly, loss of one copy of *Gdf11* decreased longevity and survival in mice, corroborating its putative role in aging. Our data demonstrate that *Gdf11* dosage is important for brain function.

## INTRODUCTION

Intellectual disability (ID) and autism spectrum disorder (ASD) affect nearly 3% of children in the United States (Zablotsky et al., 2019). These diseases can be caused by mutations in any one of hundreds of genes (Deciphering Developmental Disorders Study, 2017; Satterstrom et al., 2020), a subset of which is “dosage-sensitive”, causing neurological dysfunction by either loss-of-function mutations or an increase in expression due to copy-number gain (Rice and McLysaght, 2017). Understanding how a dosage-sensitive gene drives molecular pathogenesis can reveal the mechanisms for two diseases, the loss-of-function and increased dosage disorders (Javed et al., 2020).

Methyl-CpG binding protein 2 (*MECP2*) is the exemplary dosage-sensitive gene. *MECP2* is an X-linked gene whose loss-of-function causes Rett syndrome (RTT; OMIM: 312750) (Amir et al., 1999) and whose duplication causes *MECP2* duplication syndrome (MDS; OMIM: 300260) (Lugtenberg et al., 2009; van Esch et al., 2005). Both RTT and MDS are devastating neurological disorders characterized by intellectual disability, motor dysfunction, and seizures. Despite identification of *MECP2* as the causative gene for both disorders, we still have poor understanding of MeCP2-driven pathogenesis. MeCP2 is a methyl-cytosine binding protein that regulates gene expression in the brain (Chen et al., 2015; Gabel et al., 2015; Nan et al., 1997). Disruption of normal MeCP2 function causes subtle dysregulation of thousands of genes (Sanfeliu et al., 2019). Importantly, many of these dysregulated genes are shared between mouse models of RTT and MDS, but their expression is altered in inverse directions (Chahrour et al., 2008; Chen et al., 2015). Interrogating transcripts that are highly sensitive to MeCP2 levels can reveal disease-driving genes that could be used to develop targeted therapies for RTT and MDS (Samaco et al., 2012). Lastly, pinpointing the genes regulated by MeCP2 can enhance our understanding of molecular mechanisms of additional genes mediating neurological phenotypes (Lavery et al., 2020; Zhou et al., 2022).

To search for genes robustly regulated by MeCP2, we integrated gene expression studies that profiled RNA collected from mouse tissues bearing various *MECP2* loss- or gain-of-function alleles and discovered that Growth differentiation factor 11 (*Gdf11*) expression is highly and positively correlated with MeCP2 protein level and function. *Gdf11* is a member of the TGFβ superfamily that has been implicated in development, neurogenesis, cancer, and aging (Bajikar et al., 2017; Katsimpardi et al., 2014; McPherron et al., 1999; Wu et al., 2003); however, its role in the brain is poorly understood. Remarkably, we found that *Gdf11* expression is increased in mouse models of MDS and that genetically normalizing *Gdf11* dosage was sufficient to improve several behavioral deficits in a mouse model of MDS. Lastly, we found that loss of one copy of *Gdf11* alone is sufficient to cause broad neurological dysfunction. Our study highlights the importance of revealing key molecules downstream of dosage-sensitive genes by demonstrating that *Gdf11* is regulated by MeCP2 and that *Gdf11* dosage modifies neurological phenotypes.

## RESULTS

### MeCP2 regulates the expression of Gdf11 in the mammalian brain

Normalizing MeCP2 levels through genetic or pharmacological means is sufficient to rescue nearly all behavioral and molecular deficits in a mouse model of MDS (Sztainberg et al., 2015). For example, bolus injection of an antisense oligonucleotide (ASO) targeting *MECP2* acutely reduces MeCP2 protein in MDS mice and ameliorates behavioral deficits. We reasoned that genes whose expression was corrected after ASO treatment and whose abundance highly correlated with MeCP2 protein levels would be candidate genes to contribute to the behavioral rescue. To this end, we calculated the correlation coefficient between the fold-change of the 100 genes rescued by ASO treatment and the fold-change of MeCP2 protein levels in the hippocampi of *MECP2* duplication mice (Shao et al., 2021). We found 16 genes whose fold-change were highly correlated (Spearman’s rho > ± 0.75) with MeCP2 levels (Figure S1A). To further refine these candidates, we flagged genes that were predicted to be loss-of-function intolerant in humans as these genes could have biological consequences if their expression level is mildly altered (Karczewski et al., 2020; Lek et al., 2016). Of the 16 MeCP2-correlated genes, we identified four genes – growth differentiation factor 11 (*Gdf11*), family with sequence similarity 131 member B (*Fam131b*), veriscan (*Vcan*), and forkhead box protein P1 (*Foxp1*) with a probability of loss intolerance (pLI) greater than 0.9 (Figure 1A and Figure S1A).

**Figure 1.**
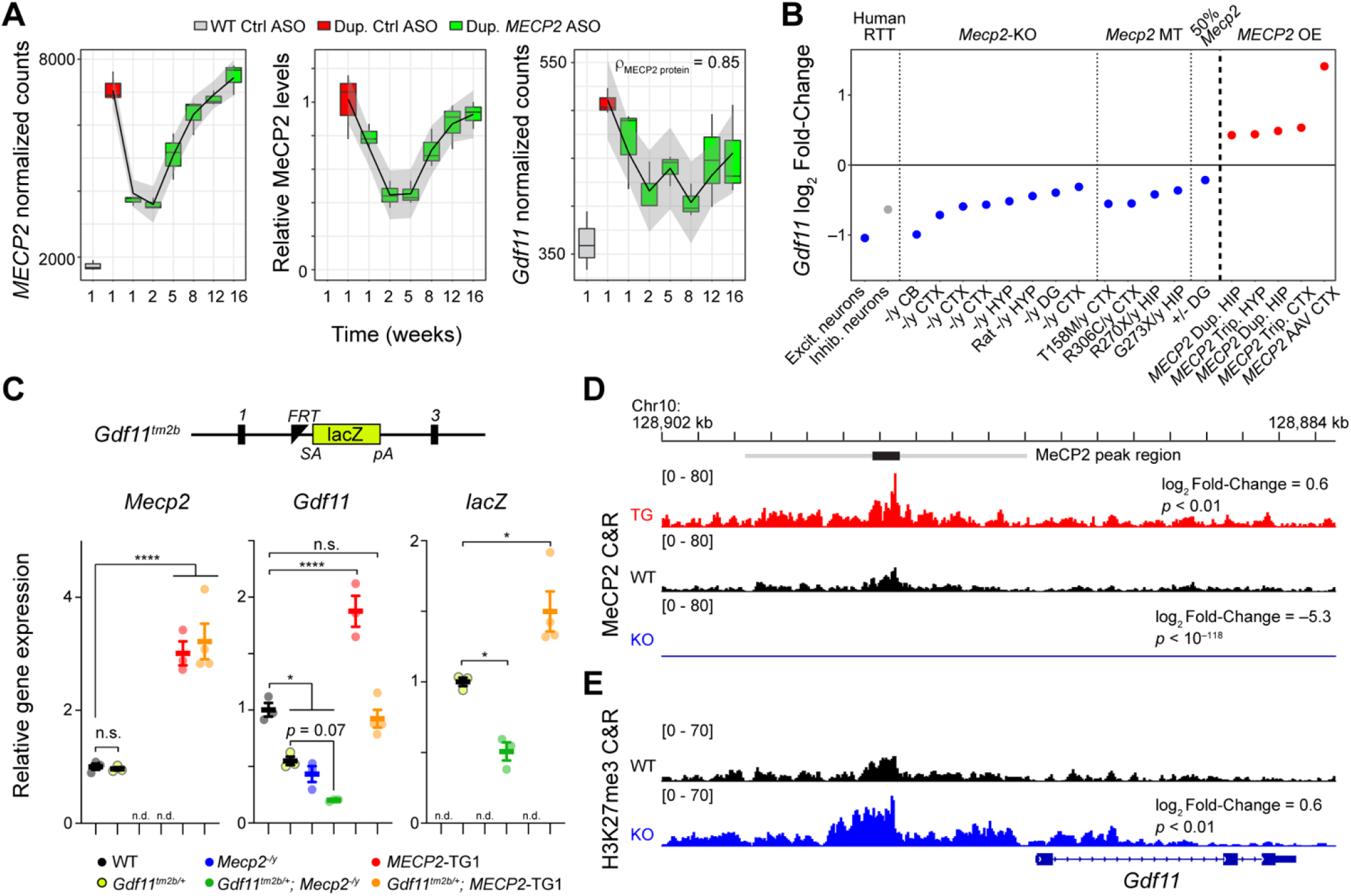
*Gdf11* is positively regulated by MeCP2. (A) Dynamic expression levels of *MECP2*, MeCP2, and *Gdf11* during anti-*MECP2* antisense oligonucleotide (ASO) treatment in *MECP2* duplication mice (data from (Shao et al., 2021)). Spearman correlation coefficient between *Gdf11* and MeCP2 is shown in the right panel. Gray interval represents a loess fit ± standard error. (B) Log_2_ fold-change of *GDF11* expression in human Rett syndrome excitatory and inhibitory neurons and *Gdf11* expression in *Mecp2*-knockout, mutant, and *MECP2* overexpression mouse model RNA-seq experiments. Grayed color indicates *p_adjusted_* > 0.1. (C) A *Gdf11-* knockout first allele (*Gdf11^tm2b^*) has a *LacZ* cassette with splice acceptor and polyA sequence knocked in between exons 1 and 3 of *Gdf11*, and exon 2 of *Gdf11* is deleted. Quantitative PCR of *Mecp2, Gdf11*, and the *LacZ* reporter from the cerebellum of wild-type, *Gdf11^tm2b/+^, Mecp2^-/y^ Gdf11^tm2b/+^; Mecp2^-/y^, MECP2-TG1*, and *Gdf11^tm2b/+^; MECP2-TG1 (n* = 3-4 biological replicates per genotype). N.d. denotes the given gene expression measurement was not detected. qPCR data is shown as mean ± sem. (D) CUT&RUN profiling of MeCP2 binding in *MECP2* transgenic mice (*MECP2*-TG1), wild-type, and *Mecp2*-knockout hippocampi at the *Gdf11* locus. Black bar represents a MeCP2 peak called by MACS2 software, and the gray bar represents an expanded region of ±3.5 kb of increased MeCP2 binding upstream of the *Gdf11* transcriptional start site. Log_2_ fold-change of MeCP2 occupancy within the gray region is shown relative to wild-type, and the *p*-value of the comparison is shown beneath the fold-change. Tracks are displayed as an aggregate of biological replicates (*n* = 3-6 biological replicates per genotype). (E) CUT&RUN profiling of H3K27me3 in wild-type and *Mecp2*-knockout at the *Gdf11* locus. Log_2_ fold-change of binding within the gray region is shown relative to wild-type, and the p-value of the comparison is shown beneath the fold-change. Tracks are displayed as an aggregate of biological replicates (*n* = 3 biological replicates per genotype). (*) *p* < 0.05, (****) *p*< 0.0001 from one way ANOVA followed by multiple comparisons testing.

Gene expression changes in *Mecp2* mutant mouse models have been inconsistent in published reports (Sanfeliu et al., 2019). To test if any of the four MeCP2-correlated, loss-intolerant transcripts were robustly sensitive to MeCP2 levels, we queried expression changes in 20 readily available transcriptional profiles generated in *MECP2* perturbed rat, mouse, and human brain samples (Supplemental Table S1). Strikingly, we found that *Gdf11* was significantly altered in 19 of the 20 profiles, whereas the other three genes did not consistently exhibit significant difference compared to control (Figure 1B and Figure S1B-D). *GDF11* levels were downregulated two-fold in human postmortem RTT excitatory neurons, while *Gdf11* levels were downregulated in the transcriptomes of *Mecp2*-null males, *Mecp2*-null heterozygous females, and a panel of *MECP2* point mutant males in multiple brain regions; these data suggest *Gdf11* is robustly sensitive to MeCP2 function. Additionally, we also found that *Gdf11* was significantly upregulated in mouse models overexpressing *MECP2* (Figure 1B). Given the strong dynamic correlation of *Gdf11* with MeCP2 levels upon ASO treatment (Figure 1A), these data suggest *Gdf11* is also sensitive to MeCP2 levels (Figure 1B). Taken together, these results demonstrate *Gdf11* is a transcript that is robustly sensitive to both MeCP2 function and protein levels.

To further investigate if MeCP2 transcriptionally regulates the *Gdf11* locus, we took advantage of a *Gdf11*-knockout allele that has a *LacZ* reporter cassette knocked into the gene (*Gdf11^tm2b^*) (Figure 1C). We crossed this allele with both a *Mecp2*-knockout (Guy et al., 2001) and a *MECP2* duplication (*MECP2*-TG1) (Collins et al., 2004) mouse model to generate a series of *Mecp2* or *Gdf11* single-mutants and *Mecp2; Gdf11* double-mutants and performed qPCR for *Mecp2, Gdf11*, and *LacZ* in the cerebellum. The expression of *Mecp2* was not changed by reduction of *Gdf11*, but *Gdf11* was significantly decreased in *Mecp2*-knockout males (*p* < 0.05) and increased in *MECP2*-TG1 animals (*p* < 0.0001). Furthermore, *Mecp2*-KO trended towards a further decrease of *Gdf11* in the *Mecp2^-/y^; Gdf11^tm2b/+^* double mutants while *Gdf11* was restored to wild-type expression in the *MECP2*-TG1; *Gdf11^tm2b/+^* double mutants. Last, expression of the *LacZ* reporter was modulated by MeCP2 levels as well, being significantly downregulated in *Mecp2*-knockout (*p* < 0.05) and upregulated in *MECP2*-TG1 animals (*p* < 0.05) (Figure 1C). These results strongly demonstrate that the *Gdf11* locus is transcriptionally sensitive to MeCP2 protein levels.

While MeCP2 has a well-described role as a transcriptional repressor and its loss leads to a derepression of gene expression, MeCP2 loss has been shown to cause a down-regulation of gene expression (Ben-Shachar et al., 2009; Boxer et al., 2020; Chahrour et al., 2008; Clemens et al., 2020; Lyst et al., 2013; Nan et al., 1998, 1997). We next sought to mechanistically understand how loss of MeCP2 leads to downregulation of *Gdf11*. We first asked if MeCP2 has enriched binding near the *Gdf11* locus. We performed <u>C</u>leavage <u>U</u>nder <u>T</u>argets & <u>R</u>elease <u>U</u>sing <u>N</u>uclease (CUT&RUN) (Skene and Henikoff, 2017) to profile MeCP2 binding in wild-type, *Mecp2*-knockout, and *MECP2-*TG1 mouse hippocampus (Figure S1E,F). We next identified peaks of enriched binding using the MACS algorithm (Zhang et al., 2008), which identified a peak upstream of the *Gdf11* transcriptional start site (TSS) (Supplemental Table S2). Due to the broad binding pattern of MeCP2 (Chen et al., 2015; Skene et al., 2010), we integrated the signal in a symmetric window straddling the peak towards the *Gdf11* TSS to robustly quantify MeCP2 occupancy. We observed a significant (*p* < 0.01) increase of MeCP2 binding upstream of the *Gdf11* TSS in the *MECP2*-TG1 hippocampus. Importantly, the MeCP2 binding signal was almost completely absent and significantly (*p* < 10^-118^) decreased in the *Mecp2*-knockout hippocampus in this same region (Figure 1D). Concurrently, we profiled the repressive histone modification H3K27me3 (Figure S1G), which has been shown to interact with and be modulated by MeCP2 (Lee et al., 2020). Within the window of MeCP2 binding upstream of the *Gdf11* TSS, we found a significant (*p* < 0.01) increase in H3K27me3 occupancy in *Mecp2*-knockout tissue and a trend towards a reciprocal decrease in H3K27me3 occupancy in *MECP2-*TG1 tissue (Figure 1E and Figure S1H). The increase in H3K27me3 in *Mecp2*-knockout tissue corroborates the downregulation we observe in *Gdf11* expression in transcriptional profiles collected from *Mecp2*-deficient models (Figure 1B). These results suggest that MeCP2 binds upstream of *Gdf11* and modifies the chromatin landscape around the *Gdf11* locus to regulate *Gdf11* expression.

### Genetic reduction of Gdf11 ameliorates behavioral deficits of MECP2 duplication mice

Since *Gdf11* dysregulation is rescued upon normalization of *MECP2* dosage, we sought to determine if normalization of *Gdf11* alone can rescue behavioral deficits in *MECP2* duplication mice. We crossed the *MECP2*-TG1 allele with the *Gdf11^tm2b^* allele to normalize *Gdf11* expression (Figure 1C). We generated a cohort of wild-type, *MECP2*-TG1, and *MECP2*-TG1; *Gdf11^tm2b/+^* double mutant littermates and behaviorally characterized these mice with an extensive battery of assays starting at sixteen weeks of age. Compared to wild-type mice, *MECP2*-TG1 mice are hypoactive when placed in an open arena, traveling less distance, and exhibiting fewer horizontal and vertical exploratory activities (Figure 2A). Normalizing *Gdf11* expression in the double mutant mice resulted in rescue of total distance traveled and horizontal activity. We also observed a partial rescue of vertical exploration and a trend towards an improvement in anxiety-like behavior in the open arena (Figure 2A and Figure S2A). Next, *MECP2*-TG1 mice experience hypoactivity and anxious behavior compared to wild-type mice when placed in an elevated plus maze apparatus, traveling less distance and entering the open arms fewer times. In contrast, we observed the double mutant animals entered the open arms and crossed the maze arms similar to wild-type animals, and the double mutant mice traveled significantly further (*p* < 0.01) on the maze compared to *MECP2*-TG1 mice (Figure 2B). Lastly, *MECP2*-TG1 mice exhibit improved learning compared to wild-type mice in a shock-tone cued fear paradigm. Specifically, mice are trained to associate a tone with a foot shock and then placed in the same environment sans tone (context) or a new environment with a tone (cue). Learning in this assay is measured by the amount of time the mouse freezes, where *MECP2*-TG1 mice freeze more in both context and cued tests. Normalization of *Gdf11* led to wild-type levels of freezing during the training routine, and normalized the cued learning compared to *MECP2*-TG1 mice (Figure S2B). Contextual learning was improved in double mutant mice with no significant difference observed compared to wild-type mice (Figure 2C). Reduction of *Gdf11* did not rescue all phenotypes present in *MECP2*-TG1 mice, as we did not observe a rescue in motor coordination on the rotating rod, change in the time spent in dark during the light-dark assay, nor any changes in sociability. However, we did observe rescue of total activity, as measured by distance traveled, in both light-dark and social assays (Figure S2C-F). These data suggest that normalization of aberrant *Gdf11* expression improves several behavioral deficits, most notably locomotion, general activity, and hippocampal learning in *MECP2*-TG1 mice.

**Figure 2.**
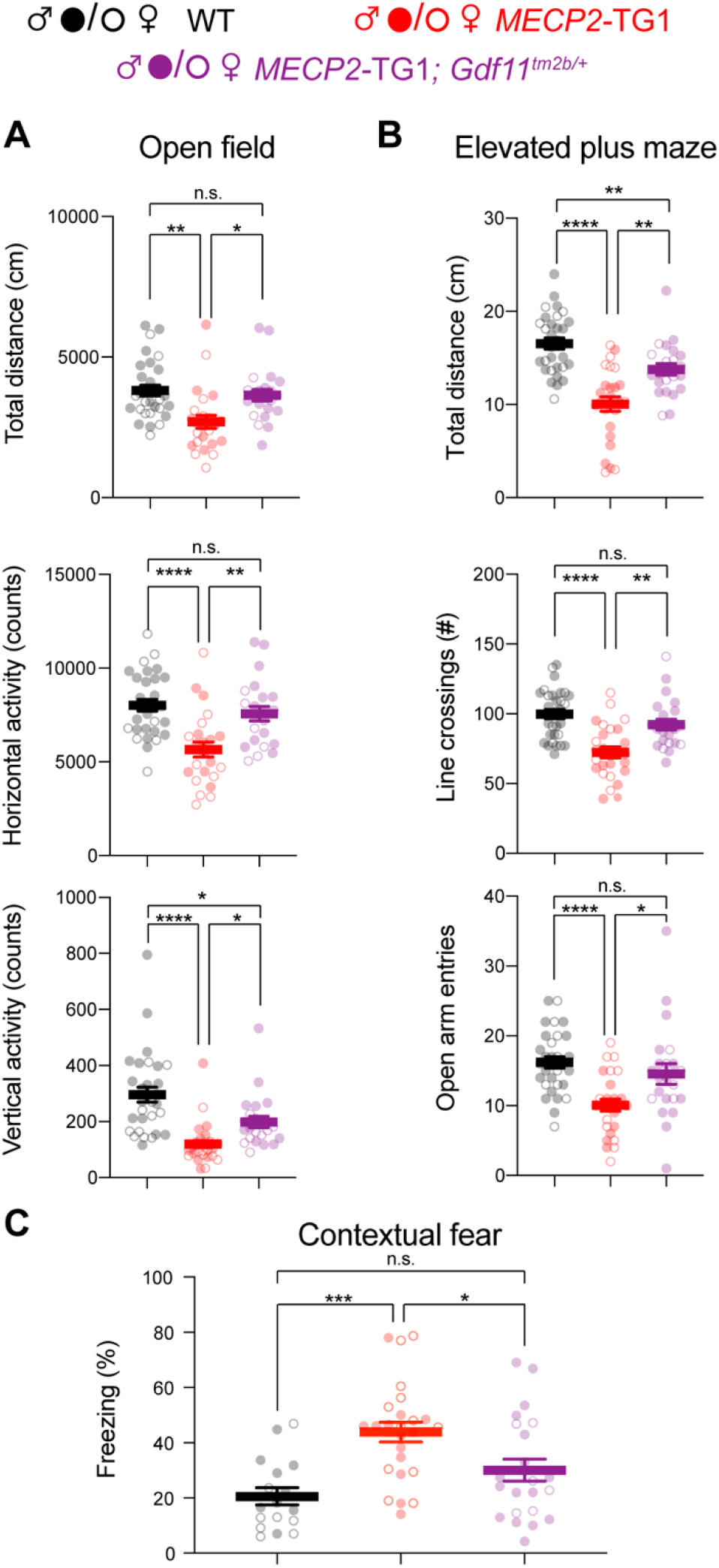
Genetic reduction and normalization of *Gdf11* dose ameliorates several behavioral deficits in *MECP2-TG1* mice. Behavioral characterization of *MECP2-TG1*, *MECP2-TG1; Gdf11^tm2b/+^* double mutants, and their respective wild-type littermate controls was performed beginning at 16 weeks of age. (A) Open field assessment of locomotion and activity. (B) Elevated plus maze assay measures of movement and anxiety. (C) Learning assessment using contextual fear-conditioning. Greater freezing indicates better memory of the context. Central estimate of data is shown as mean ± sem. Closed circles denote male mouse data points and open circles denote female mouse data points. For all measurements, *n* = 30 wild-type mice (19 male, 11 female); *n* = 24 *MECP2-TG1* mice (12 male, 12 female); and *n* = 22 *MECP2-TG1; Gdf11^tm2b/+^* mice (16 male, 6 female). All data were analyzed using a Welsch one way ANOVA with Dunnett’s post hoc multiple comparisons. * *P* < 0.05, ** *P* < 0.01, *** *P* < 0.001, and **** *P* < 0.0001.

### Loss of one copy of Gdf11 causes neurobehavioral deficits in mice

The previous results raised the possibility that mouse neurological function is sensitive to changes in *Gdf11* dosage. Supporting this evidence, loss of one copy of *GDF11* function causes a developmental disorder with neurological features in humans (Ravenscroft et al., 2021). Furthermore, *GDF11* is broadly and highly expressed in both the human and mouse brains, suggesting a potentially important involvement role in brain function (Figure S3A,B) (Li et al., 2017).

To determine if loss of one copy of *Gdf11* resulted in neurobehavioral deficits, we generated a cohort of *Gdf11^tm2b/+^* mice to characterize with a comprehensive battery of tests. We first confirmed that *Gdf11^tm2b/tm2b^* mice exhibited abnormal development and perinatal lethality as previously described (McPherron et al., 1999), which along with a 50% reduction of *Gdf11* in the brains of *Gdf11^tm2b/+^* mice (Figure 1C), demonstrating that the *Gdf11^tm2b^* allele is a null allele (Figure S3C). Thus, *Gdf11^tm2b^* heterozygotes can be used to model the effects of losing one copy of *Gdf11*. Given these results, we generated a cohort of wild-type and *Gdf11^tm2b/+^* littermates and began general health assessments at weaning. We observed that these mice did not display any overt phenotypes and did not have a change in body weight in either sex through till sixteen weeks of age (Figure S3D).

We began deeper behavioral characterization of the cohort at sixteen weeks of age. In the open field assay, we observed that *Gdf11^tm2b/+^* mice are hyperactive as indicated by a significant increase in total distance traveled (*p* < 0.05), a significant increase in horizontal activity counts (*p* < 0.05), and a significant increase in vertical exploratory activity counts (*p* < 0.01) (Figure 3A). *Gdf11^tm2b/+^* mice explored the center of the arena at a similar rate as wild-type mice (Figure S3E). In the elevated plus maze, *Gdf11^tm2b/+^* mice also displayed hyperactivity, traveling a significantly further distance (*p* < 0.01) and decreased anxiety, entering the open arm a significantly higher number of times (*p* < 0.001) (Figure 3B). *Gdf11^tm2b/+^* mice traveled a significantly further distance (*p* < 0.05) in the light-dark assay while spending significantly more time (*p* < 0.05) in the dark (Figure S3F). Furthermore, when placed in a 3-chamber apparatus, *Gdf11^tm2b/+^* mice explored more both during habituation and testing phases (Figure 3C and Figure S3G). This manifests with a significantly (*p* < 0.05) decreased time socializing compared to control, though the mice still prefer interacting with the partner mouse over the object. The ratio of time spent investigating the partner mouse over the total investigation time was significantly (*p* < 0.0001) less in *Gdf11^tm2b/+^* mice (Figure S3H). Next, we observed a significant (*p* < 0.0001) reduction in motor coordination in learning as assessed by the rotating rod assay (Figure 3D). Finally, we observed a trend in reduction in learning during the training phase of a shock-tone conditioning paradigm (Figure S3I), while observing a significant reduction in contextual learning (*p* < 0.01) and cued learning (*p* < 0.05) (Figure 3E). These data collectively suggest that neurological behaviors are sensitive to loss of one copy of *Gdf11* in mice.

**Figure 3.**
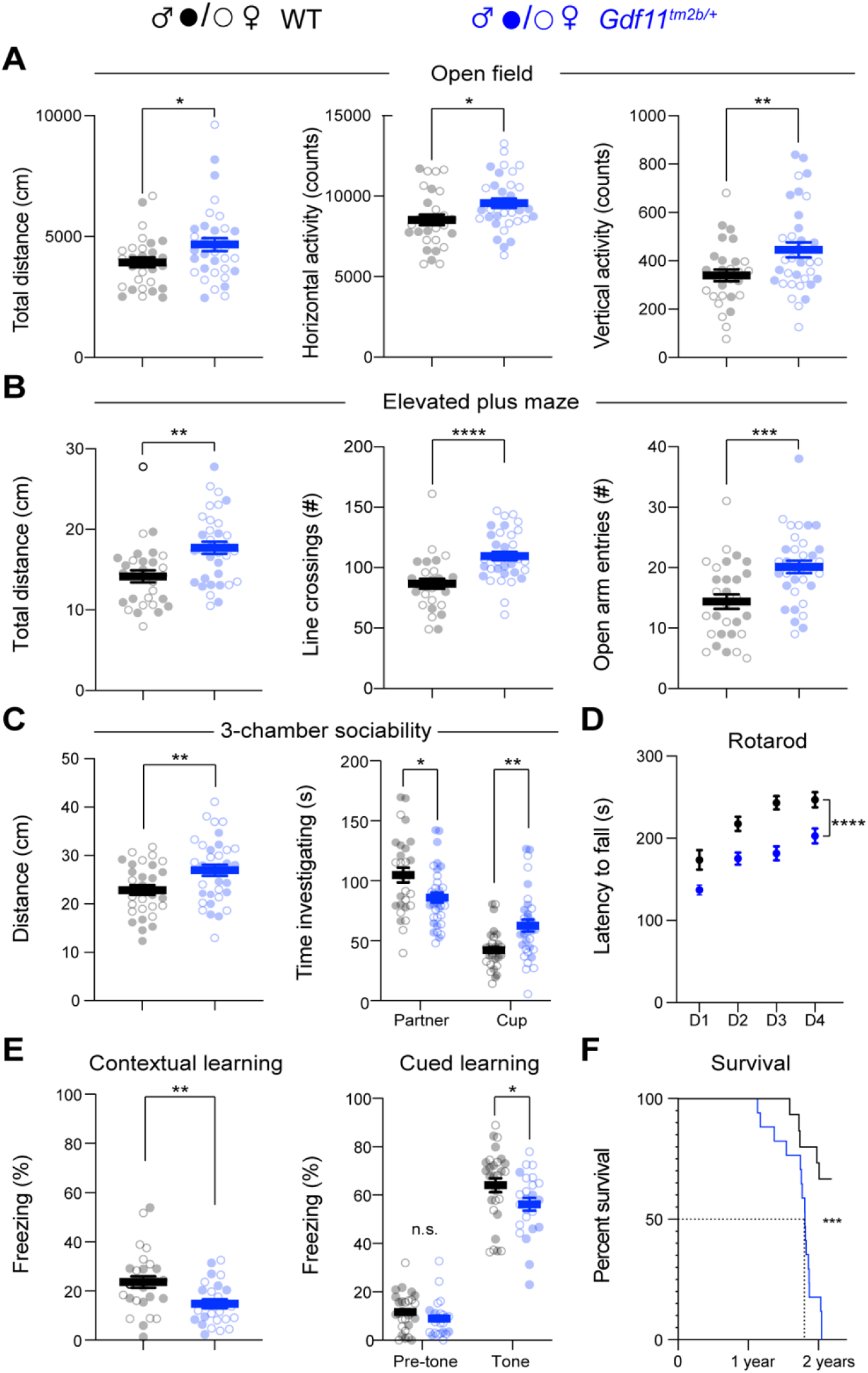
Mice lacking one copy of *Gdf11* (*Gdf11^tm2b/+^*) display behavioral deficits. Behavioral characterization of *Gdf11^tm2b/+^* mice and their respective wild-type littermate controls was performed beginning at 16 weeks of age. (A) Open field assessment of locomotion and activity. (B) Elevated plus maze assay measures of movement and anxiety. (C) Distance traveled and time investigating novel mouse in 3-chamber sociability assay. (D) Motor coordination measured by the rotating rod assay. (E) Learning assessment using contextual and cued fear-conditioning. Greater freezing indicates better memory of the context. (F) Survival analysis. Central estimate of data is shown as mean ± sem. Closed circles denote male mouse data points and open circles denote female mouse data points. For behavioral measurements except fear conditioning, *n* = 29 wild-type mice (15 male, 14 female) and *n* = 34 *Gdf11^tm2b/+^* mice (16 male, 18 female). For fear conditioning assay (I), *n* = 28 wild-type mice (16 male, 12 female) and *n* = 24 *Gdf11^tm2b/+^* mice (9 male, 15 female). For survival analysis, *n* = 15 wild-type mice and *n* = 17 *Gdf11^tm2b/+^* mice. Behavioral data were analyzed using Welsch t-test. Survival was analyzed using Mantel-Cox test. * *P* < 0.05, ** *P* < 0.01, *** *P* < 0.001, and **** *P* < 0.0001.

As *Gdf11* has been implicated to naturally decrease in aging in mice (Poggioli et al., 2016), we tested if a reduction in *Gdf11* dosage can alter aging by tracking the survival of a cohort of *Gdf11^tm2b/+^* mice. Strikingly, we found a significant (*p* < 0.001) reduction in survival of *Gdf11^tm2b/+^* mice compared to wild-type littermates (*n* = 15 wild-type and *n* = 17 *Gdf11^tm2b/+^*), exhibiting a median survival of 1.8 years (Figure 3F). Also, a portion *Gdf11^tm2b/+^* mice were euthanized due to reaching ethical endpoints including poor body conditioning, self-lesioning, and development of tumors, which suggests that *Gdf11* dosage plays an important role in long-term health and survival in mice.

### Loss of one copy of Gdf11 does not alter proliferation, amplifying progenitors, or neural stem cells in the adult hippocampus

Learning and memory is linked to the generation of adult-born neurons in the subgranular zone (SGZ) of the mouse dentate gyrus (Kempermann and Gage, 2002). Complete loss of *Gdf11* in adulthood was shown to increase proliferation in the SGZ, specifically in the number of amplifying progenitors (SOX2 positive), while impairing the overall number of newborn neurons that survive and integrate into the existing hippocampal circuitry (Mayweather et al., 2021). Additionally, neurogenesis is negatively impacted in the olfactory epithelium with loss or inhibition of GDF11 (Wu et al., 2003). To test if loss of one copy of *Gdf11* altered overall proliferation in the SGZ, potentially acting as a mechanism for the changes in learning and memory observed in *Gdf11^tm2b/+^* mice, we labeled newborn, dividing cells by injecting *Gdf11^tm2b/+^* and control mice with 5-Ethyl-2’-deoxyuridine (EdU). We injected the animals with EdU (10 mg/kg) for 5 consecutive days and fixed the brains at 7 days after the first injection. We performed these injections in sixteen-week-old animals, matching the timepoint when we performed behavioral characterization. We then stained for EdU, SOX2, and GFAP to label proliferating cells (EdU^+^), neural stem cells (SOX2^+^GFAP^+^) and amplifying progenitors (SOX2^+^) (Zhao and van Praag, 2020). We did not observe any significant changes in the number or density of these cell populations in *Gdf11^tm2b/+^* mice using quantitative stereology (Figure 4 and Figure S4A-D). Furthermore, no gross changes in brain anatomy or volume of the dentate gyrus were observed in *Gdf11^tm2b/+^* mice (Figure S4E-G). These results indicate that the behavioral deficits caused by loss of one copy of *Gdf11* are not due to altered proliferation in the adult mouse SGZ.

**Figure 4.**
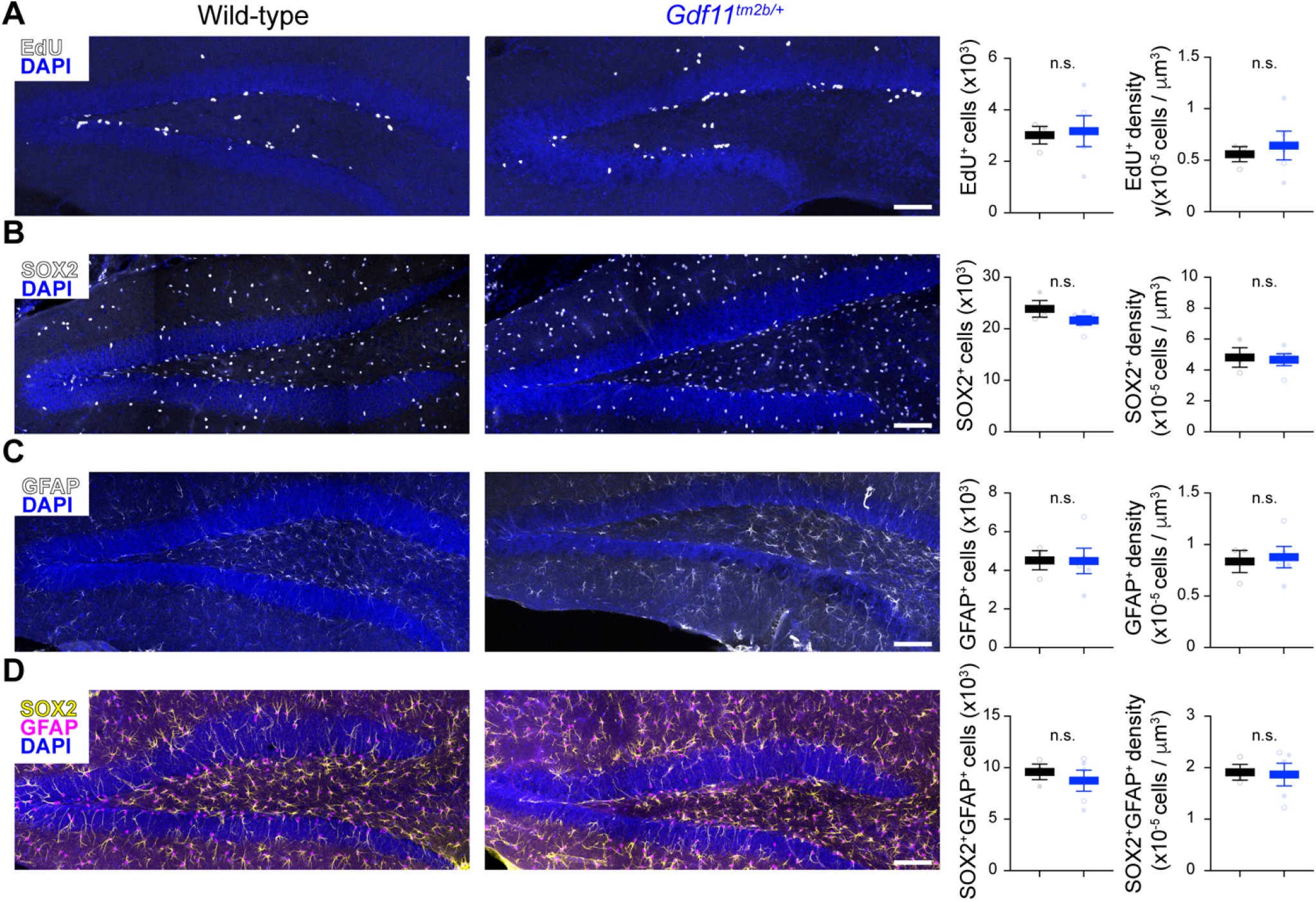
Loss of one copy of *Gdf11* does not alter adult neurogenesis in the dentate gyrus. Quantification of markers of (A) proliferative cells (EdU) or (B-D) neural progenitor pools (SOX2 and GFAP) in the subgranular zone of the dentate gyrus in wild-type or *Gdf11^tm2b/+^* mice. Representative images of the dentate gyrus for the indicate stains are shown on the left. The projected total number of cells and the density of cells is shown on the plots to the right using stereology to quantify *n* = 3-6 separate slices per animal (*N* = 3 wild-type animals and 5 *Gdf11^tm2b/+^* animals) (see Methods). Each data point is the aggregated value for one animal. Closed circles denote male animals and open circles denote female animals. Data are presented as mean ± sem. Data were analyzed using Welsch’s t-test. Scale bar is 200 μm.

## DISCUSSION

The heterogeneity in genetic causes for IDD and ASD has precluded the development of targeted therapies for these disorders. Distilling any common molecular processes that have gone awry in multiple disorders would give us a handle to investigate further and identify drug targets. In this study, we thoroughly investigated genes regulated by the exemplar dosage-sensitive gene *MECP2*, the causative gene of two different neurological disorders (RTT and MDS) to discover that MeCP2 sensitively and acutely regulates *Gdf11*. Intriguingly, *Gdf11* is inversely dysregulated in RTT and MDS and restoring normal *Gdf11* dosage ameliorated several behavioral deficits in a mouse model of MDS. This potential sensitivity of *Gdf11* dosage in the brain led us to further discover that reduction of *Gdf11* dosage alone caused neurobehavioral deficits in mice as well. Our study strongly suggests that GDF11 signaling is dysregulated in more than one neurodevelopmental disorder (RTT, MDS, and *GDF11* haploinsufficiency) and modifies MeCP2-driven abnormal behaviors.

Transcriptomics is often used to identify molecular pathways that are dysregulated in diseases. However, it can be challenging to uncover the core genes that are altered can be challenging due to secondary effects and neuronal dysfunction. These challenges have been particularly problematic in using transcriptomics to dissect the molecular pathogenesis of MeCP2 (Sanfeliu et al., 2019). In this study, we leveraged dynamic gene expression changes upon acute decrease of *MECP2* in a mouse model of MDS (Shao et al., 2021), which specifically revealed that *Gdf11* is highly correlated with MeCP2 protein levels. Generalization of these results demonstrated that *Gdf11* is robustly dysregulated in mouse models of MeCP2-related disorders and regulated by MeCP2 at the chromatin level (Figure 1). These data suggest that *Gdf11* is proximally downstream of MeCP2, sensitive to both MeCP2 expression and function. Ongoing efforts are looking to restore MeCP2 dosage in mouse models of RTT and MDS (Shao et al., 2021; Sinnamon et al., 2017; Sinnett et al., 2021). Typically, these studies rely on behavioral outcomes to assess treatment efficacy. However, behavioral improvements downstream of restored MeCP2 dosage or function may take weeks and may require a large sample size to manifest (Katz et al., 2012; Shao et al., 2021; Sztainberg et al., 2015). We propose that *Gdf11* expression in the brain can be used as an endogenous reporter for MeCP2 dosage and function in preclinical studies that can leverage the sensitivity of biomolecular measurements. Adding *Gdf11* to the repertoire of metrics to evaluate treatment outcomes may prevent preliminary and promising candidates from being scrapped if behavioral outcomes are unclear or slow to normalize.

We showed that normalization of *Gdf11* dosage improved several behavioral deficits in a mouse model of MDS when normalization of *Gdf11* occurred using a constitutive germline deletion. If normalization of *Gdf11* dosage could be used to treat MeCP2-related disorders, we must first test if restoring *Gdf11* dosage in symptomatic RTT or MDS mouse models leads to outcome improvements. These experiments will decouple if MeCP2 regulation of *Gdf11* is critical during a presymptomatic window or throughout life. Importantly, GDF11 is a secreted factor, and its protein dosage could be modulated using recombinant protein to increase dosage (for RTT or *GDF11* haploinsufficiency) or using a neutralizing antibody to decrease dosage (for MDS) (Morvan et al., 2017; Ozek et al., 2018; Zhang et al., 2018).

Our study contributes to the growing body of work that *GDF11* is a dosage sensitive gene important to neural function. Loss-of-function mutations in *GDF11* have recently been described to cause neurological dysfunction as part of a broader developmental disorder. These patients exhibit abnormalities including intellectual disability, speech delay, and seizures (Cox et al., 2019; Ravenscroft et al., 2021). In this study, we show that reducing *Gdf11* dosage in mice also causes abnormal phenotypes, which we can now leverage to investigate mechanisms of neuronal dysfunction in *GDF11* haploinsufficiency. Furthermore, a recent genomic study predicted that *GDF11* is intolerant to copy number gains in humans (Collins et al., 2022). Our work suggests that increased *Gdf11* expression contributes to abnormal phenotypes in mouse models of MDS. Using mouse models that overexpress *GDF11* alone could be used to determine if copy number gain in GDF11 is toxic (Jones et al., 2018). These studies will be important to determine whether GDF11 can be a therapeutic target if appropriately dosed.

Though *GDF11* is broadly and highly expressed in the mouse and human brains, we have a limited understanding of its function. As a secreted factor, it is unclear why this gene must be expressed throughout the brain in multiple cell types (Saunders et al., 2018). This may suggest that GDF11 has a strong autocrine and paracrine role in the brain rather than an endocrine role. We must therefore catalog which neurons or brain regions secrete or sense GDF11 and understand the functional neuronal response to GDF11 stimulation. These studies will parse out the specific relationship of GDF11 dosage and neurological phenotypes. For example, we see robust hyperactivity in *Gdf11^tm2b^* heterozygotes, but more variable effects on anxiety. Is this influenced because specific brain regions or cell types are producing or sensing GDF11? Furthermore, we discovered that *Gdf11* dosage is important to longevity and survival in mice, corroborating a role of GDF11 in aging (Katsimpardi et al., 2014; Loffredo et al., 2013; Sinha et al., 2014), but is this role brain-specific? As such, understanding the production, transduction, and transmission mechanisms of GDF11 signaling will have long standing importance to multiple fields.

In summary, our study provides a framework for identifying putative disease-modifying genes by integrating multiple -omics level datasets. Through these analyses, we linked seemingly disparate disease-causing genes, *MECP2* and *GDF11* in this case, by characterizing a regulatory relationship between them. Probing other disease-causing genes in this way will allow us to stitch together a network of dosage-sensitive genes, which will reveal points of regulatory and molecular convergence across a wide range of diseases.

## MATERIALS AND METHODS

### Animals

Baylor College of Medicine Institutional Animal Care and Use Committee (IACUC, Protocol AN-1013) approved all mouse care and manipulation. Mice were housed in an AAALAS-certified level 3 facility on a 14-hour light cycle. Sperm from *Gdf11^tm2a(EUCOMM)Hmgu^* mice was obtained from the MMRRC (Stock ID: 037721-UCD, RRID: MMRRC_037731-UCD). In vitro fertilization was performed with C57BL6/J donor eggs by the Genetic Engineering Mouse Core at Baylor College of Medicine. *Gdf11^tm2a^* mice were maintained on C57BL6/J background. *Gdf11^tm2b^* mice were generated by crossing male *Gdf11^tm2a^* mice with *Sox2-Cre* female mice obtained from Jackson lab (Stock No: 008454; RRID: IMSR_JAX:008454). *Gdf11^tm2b^* mice were maintained on C57BL6/J background. *MECP2-TG1* and *Mecp2-KO* mice were previously described (Collins et al., 2004; Guy et al., 2001), maintained on a C57BL6/J background, and available from Jackson lab (Stock Nos: 008679; RRID: IMSR_JAX:008679 and 003890; RRID: IMSR_JAX:003890). *Gdf11* and *MECP2* double mutant mice were generated by breeding male *Gdf11^tm2b/+^* mice to female *MECP2-TG1* or *Mecp2^+/-^* mice. Mice were monitored daily by veterinary staff. Survival age was taken as the age in which a mouse died of natural causes or when a mouse was euthanized due to poor body condition, self-lesioning, or tumor growth according to the institutional veterinarian guidelines. All procedures to maintain and use these mice were approved by the Institutional Animal Care and Use Committee for Baylor College of Medicine and Affiliates.

### RNA-seq meta-analysis

#### Analysis of transcripts correlated with MeCP2 protein levels

To identify genes that correlate with MeCP2 protein levels, we reanalyzed the transcriptional response of humanized *MECP2* duplication mice treated with an anti-*MECP2* ASO over time (GSE #1512222). The differential gene expression was perform as previously described (Shao et al., 2021), and the log_2_ fold-changes of the genes significantly altered between control and *MECP2* duplication mice per time point relative to control were extracted. The log_2_ fold-change in MeCP2 protein levels between control-treatment and *MECP2*-ASO treatment for a similar cohort of treated mice was taken from Fig. 4C of that paper. The Spearman correlation was calculated between protein level fold-change and gene expression level foldchange. The probability of loss intolerance (pLI) was extracted from the gnomad data base (Karczewski et al., 2020).

#### Extraction of gene expression changes from MeCP2 related transcriptional profiles

To extract the expression of RNA fold-changes for candidate, correlated genes across MeCP2-perturbed, we extracted log_2_ fold-change of each transcript (MeCP2 mutant allele / wild-type) and an associated *p*_adjusted_. A gene was considered significant if the false-discovery rate was below 0.1. Bulk RNA-sequencing data from MeCP2-perturbed models were preferred but in three unique cases: 1) Single-cell data from postmortem RTT tissue (Renthal et al., 2018), microarrays from two *MECP2* truncation models that do not have a corresponding RNA-sequencing data set (Baker et al., 2013), and bulk nuclei RNA-seq from *MECP2* AAV expressing neurons that do not have a corresponding bulk RNA-sequencing data set (Ito-Ishida et al., 2020). Data were extracted from Supplemental Tables or processed files deposited on GEO as possible. An additional set of data tables were found in the source data from a previous meta-analysis published in (Raman et al., 2018). The remaining data tables were generated by obtaining raw counts from GEO and re-analyzing using DESeq2 between MeCP2 perturbed samples and wild-type samples. A summary of the studies and the sources of data tables are shown in Supplemental Table S1.

### CUT&RUN epigenetic profiling

#### Nuclear isolation

Nuclei were isolated from frozen 8-week-old hippocampi from *MECP2*-TG1, *Mecp2-KO*, and wild-type littermates (*n =* 3 for *MECP2*-TG1 and *Mecp2*-KO, *n* = 6 for wild-type) using an iodixanol gradient modified from (Mo et al., 2015). Briefly, both flash frozen hippocampi per animal were dropped into a 7 ml dounce homogenizer containing 5ml buffer HB (0.25M Sucrose, 25mM KCl, 2mM Tricine KOH pH 7.8, 500uM Spermidine) and dounced 10x with loose pestle A and 20x with tight pestle B. Then 320μl of HB-IGEPAL (HB Buffer + 5% IGEPAL CA-630) was added to each homogenizer and dounced 20x more with tight pestle. Each sample was incubated for 10 mins on ice and filtered through a 30μM filter into a conical tube containing 5ml of iodixanol working solution (5 volumes Optiprep (Sigma Aldrich, D15556) + 1 volume Optiprep Diluent (150mM KCl, 30mM MgCl_2_, 120mM Tricine-KOH pH 7.8)) and mixed by tube inversion.

To set up the gradient, 4ml of 40% Iodixanol (3 volumes working solution + 1 volumes HB buffer) was added to a 50ml round bottomed conical tube. Then, 7.5 ml 30% Iodixanol (3 volumes working solution + 2 volumes HB) was slowly overlayed on top, followed by 10ml of the sample containing mixture prepared above. This gradient was spun at 10,000g for 20 mins at 4°C in a hanging bucket centrifuge (Sorvall Lynx 6000) with “decel” turned off. After centrifugation, nuclei are located at the interface between 30% and 40% iodixanol layers. Iodixanol containing supernatant above the nuclei was slowly discarded with bulb pipette. Approximately 1.5-2ml of the interface containing nuclei were collected and placed into 2ml microcentrifuge tube.

The number of nuclei were quantified by taking 20μl of sample and mixing it with 2μl 0.2mg/ml DAPI diluted in HB buffer. After three-minute incubation at RT, the nuclei were diluted 1:10 in buffer HB and counted on a Countess II with DAPI channel to quantify.

#### CUT&RUN

Cleavage Under Targets & Release Nuclease (CUT&RUN) was performed on nuclei isolated from above following (Skene and Henikoff, 2017). Briefly, we performed one nuclear isolation per animal and then split the nuclei into three individual tubes for the three antibodies surveyed (MeCP2, H3K27me3 and IgG). An individual sample will refer to one antibody from a unique animal.

To activate Concanavalin A coated magnetic beads (Bangs Laboratories Inc., #BP531) for binding, we incubated 25μl per sample with 3x volumes binding buffer (20mM HEPES-NaOH 7.5, 10mM KCl, 1mM CaCl_2_, 1mM MnCl2), rotated at RT for 5 minutes, and washed 2x with 1ml binding buffer. All washes are done by placing microcentrifuge tube on magnetic rack and waiting until solution is clear as beads separated from the solution. Following washes, the beads were resuspended in 50μl binding buffer per sample.

After bead activation, 200μl beads were added to 1.5×10^6^ nuclei in the iodixanol mixture from each animal and rotated for 10 minutes at room temperature. Following binding of nuclei to beads, all further processing was done on ice. Bead bound nuclei from each animal was washed 2x with 1.5 ml wash buffer (20mM HEPES NaOH pH 7.5, 150mM NaCl, 500uM Spermidine (Sigma #S0266), and 0.5% Ultrapure BSA (Invitrogen #AM2618) with 1 tablet of Complete Protease Inhibitor Cocktail (Roche #11873580001) per 50ml). After the second wash, 250 μl of nuclei bound beads (~250,000 nuclei) in wash buffer were added to individual microcentrifuge tubes corresponding to each antibody.

Supernatant was removed and beads were resuspended in 250μl of antibody buffer (wash buffer + 0.05% Digitonin (Calbiochem #11024-24-1) + 2mM EDTA) containing an individual antibody – rabbit anti-MeCP2 (1:100, Cell Signaling #3456, clone D4F3, RRID: AB_2143894), rabbit anti-Tri-Methyl-Histone H3 (Lys27) [H3K27me3] (1:100, Cell Signaling #9733, clone C36B11. RRID: AB_2616029), and rabbit IgG (1:100, Millipore #12-370, RRID: AB_145841). Antibody buffer was added to nuclei bound beads during light vortexing (1100 rpm). Tubes were then placed at 4°C to rotate overnight. Following rotation, a quick spin on a microcentrifuge was performed to remove liquid from the cap and then washed 2x with 1ml Dig-wash buffer (wash buffer + 0.05% digitonin). Following the second wash, samples were resuspended in 200μl dig wash buffer and transferred to PCR strip tubes. Supernatant was removed and beads were resuspended in 100μl 1x pAG-MNase (Epicypher #15-1016); 20x stock pAG-MNase) in dig-wash buffer and mixed with gentle flicking. Tubes were placed on nutator for 1hr at 4°C. Following incubation, samples were washed with 200μl dig-wash buffer, transferred to new 1.5ml microcentrifuge tube, and washed 1 additional time in 1 mL dig-wash buffer.

To initiate cleavage and release of DNA bound fragments, each sample was resuspended in 150μl of dig-wash buffer while gently vortexing. Samples were placed on ice in 4°C room for 10 minutes to equilibrate. To start digestion, while in 4°C room, 3μl of 100μM CaCl_2_ was added to each tube, quickly flicked, and immediately returned to ice. Following 45-minute incubation on ice, 150μl 2x STOP (340mM NaCl, 20mM EDTA, 4mM EGTA, 0.05% Digitonin, 100μg/ml RNAse A (ThermoFisher Scientific #EN0531), 50 μg/ml Glycogen (ThermoFisher Scientific #10814010) and 1 ng E. coli spike-in / sample (Epicypher #18-1401) mixture was added to each sample. Tubes were then incubated at 37°C for 30 mins to digest RNA and release DNA. E.coli spike-in control was used for normalization of CUT&RUN signal due to differences in library amplification and/or sequencing. Supernatant was transferred to new tube and incubated with 1.5μl 20% SDS and 5μl 10mg/ml Proteinase K while lightly shaking at 50°C for 1hr. DNA was then purified by phenol-chloroform extraction using Maxtract Tubes (129046, Qiagen) and pellet resuspended in 36.5 μl TE Buffer.

#### CUT&RUN Next Generation Sequencing Library Preparation

Library preparation was modified from protocols.IO (dx.doi.org/10.17504/protocols.io.bagaibse) utilizing reagents from the NEB Next II DNA Ultra Kit (New England Biolabs #E7645S) and Unique Combinatorial Dual index kit (New England Biolabs #E6442S,) with modifications outlined below. Input DNA was quantified with Qubit, and 6 ng of CUT&RUN DNA was used as input for the H3K27me3 samples and 25μl of CUT&RUN DNA was used for both MeCP2 and IgG samples. Volume of DNA was brought up to 25ul and 1.5μl End Prep Enzyme Mixture and 3.5 μl Reaction buffer were added and incubated at 20°C for 30 mins and 50°C for 60 mins. After end prep, 15 μl of NEB Next Ultra Ligation Mastermix, 0.5 μl Ligation Enhancer and 1.25 μl of Adapter (1.2 pmol adapter for H3K27me3, 0.6 pmol adapter for MeCP2 and IgG) were added directly to the PCR tube, mixed by pipetting, and incubated for 15 mins at 20°C. Then, 1.5μl of USER Enzyme is added to each tube. Finally, SPRI select beads (Beckman Coulter # B23318) were used at 1.6x ratio to remove excess adapter and eluted in 15μl of TE buffer.

PCR amplification was performed using 13μl of adaptor ligated fragments, 1μl of Unique Combinatorial Dual Index (one index per sample), 1μl of sterile water, and 15μl 2x Q5 Master Mix. 14 cycles of PCR were performed with 10 seconds of denaturation at 98°C and 10 sec of annealing/extension at 65°C. Following PCR amplification, SPRI select beads were used for two-sided size selection; 0.65x right sided selection was performed first followed by 1.2x left sided size selection. Sample was eluted in 15μl TE.

For quality control, each library size distribution was determined by Agilent Tapestation HS DNA 1000 (Agilent Technologies #5067) and concentration was determined by KAPA PCR (Roche #07960140001). Libraries were pooled together at equimolar concentrations and submitted to the Baylor Genomic and RNA Profiling Core. Each library was sequenced for approximately 40 million paired end reads of 100bp in length on a Novaseq S1 flow cells.

### CUT&RUN analysis

#### CUT&RUN Sequence Alignment

Our CUT&RUN data analysis pipeline was adapted from CUT&RUN Tools (Zhu et al., 2019). Raw Fastq files were appended together using Linux cat function. Adapter sequences were removed from sequence reads using Trimmomatic version 0.36 (2:15:4:4:true LEADING:20 TRAILING:20 SLIDINGWINDOW:4:15 MINLEN:25) from the Truseq3.PE.fa adapter library and kseq (Bolger et al., 2014; Zhu et al., 2019).

Confirmation of adapter removal and read quality was performed with fastqc (v0.11.8). Alignment was performed with bowtie2-2.3.4.1 (--dovetail --phred33) to both mm10 (GENCODE GRCm38p6 primary assembly version 18) and the spike-in Ecoli K12 Genomes (GCF_000005845.2_ASM584v2). Bedtools (v2.29.1) was used to process BAM files to BED files, remove blacklist (mm9 blacklist lifted over to mm10 and combined with mm10 blacklist downloaded with CUT&RUNTools (Zhu et al., 2019) and to generate bedgraphs (Quinlan and Hall, 2010). Each sample was normalized to internal Ecoli spike-in utilizing spike_in_calibration.sh as described previously (Meers et al., 2019). Both spike-in normalized bedgraphs for each sample and merged bedgraphs were converted to bigwigs using UCSC bedgraph to bigwig. Spike-in normalized bigwigs from each genotype were merged using deeptools bigwigCompare and averaged for summary figures.

#### MeCP2 Peak Calling

Peaks were called using the MACSr package (version 1.0.0) using the following parameters: gsize = “mm”, format = “BAMPE”, broad = “TRUE”, keepduplicates = “all”, qvalue = 0.0001 (Zhang et al., 2008). Input to the peak calling function callpeak were the six replicates of wild-type MeCP2 BAM files and three replicate *Mecp2-* KO MeCP2 BAM files as control. Peaks with a q-value less than 0.0001 were retained for further analysis. Peaks were associated with the closest gene using bedtools -closest and mm10 GRCm38 gene coordinates. Visualization and Quality Control of CUT&RUN signal

Integrated Genome Viewer (IGV) v2.11.1 was used to examine spike-in normalized bigwig tracks at individual loci (Robinson et al., 2011). We extracted the integrated density at MeCP2 peaks ±3.5 kb of each filtered peak by using the bedtools function multicov for each BAM file generated (MeCP2, H3K27me3, and IgG). We performed differential binding using the DESeq2 package (version 1.32) using the E-coli spike in control counts as normalization factor and cooksCutoff set to FALSE. DESeq2 was used to perform principal component analysis of each mark individually and then together. Quantification of MeCP2 binding was performed across all samples, pooling two batches of sample collection due to no apparent batch effects by PCA. Quantification of H3K27me3 binding was performed in two batches, with *MECP2*-TG1 and *Mecp2-KO* samples assessed with their respective wild-type samples within the batch.

### RNA extraction, reverse transcription, and qPCR

Total RNA was isolated from one cerebellar hemisphere using the Qiagen miRNeasy Mini kit (Qiagen #217004). On column DNAse digestion (Qiagen #79254) was performed according to manufacturer’s protocol to remove genomic DNA. 2 μg of total RNA was used to synthesized cDNA using the M-MLV reverse transcriptase (Life Technologies) according to manufacturer’s protocol. qRT-PCR was performed using a CFX96 Real-Time System (Bio-Rad) using PowerUp SYBR Green Master Mix (ThermoFisher #A25741), 0.4 uM forward and reverse primers, and 1:20 dilution of cDNA. The following cycling conditions were used: 95°C for 5 min, 39 cycles of 95°C for 11 sec, 60°C for 45 sec, plate read, a final melt of 95°C, and melt curve of 65-95°C at +0.5°C increments. The specificity of the amplification products was verified using melt-curve analysis. The Ct values were calculated with the Bio-Rad software, and relative gene expression was calculated using the ΔΔCt method using *Ppia* for normalization. All reactions were performed in technical duplicate with a minimum of three biological replicates. Data are presented as mean ± SEM in main figure.

#### qPCR primer sequences

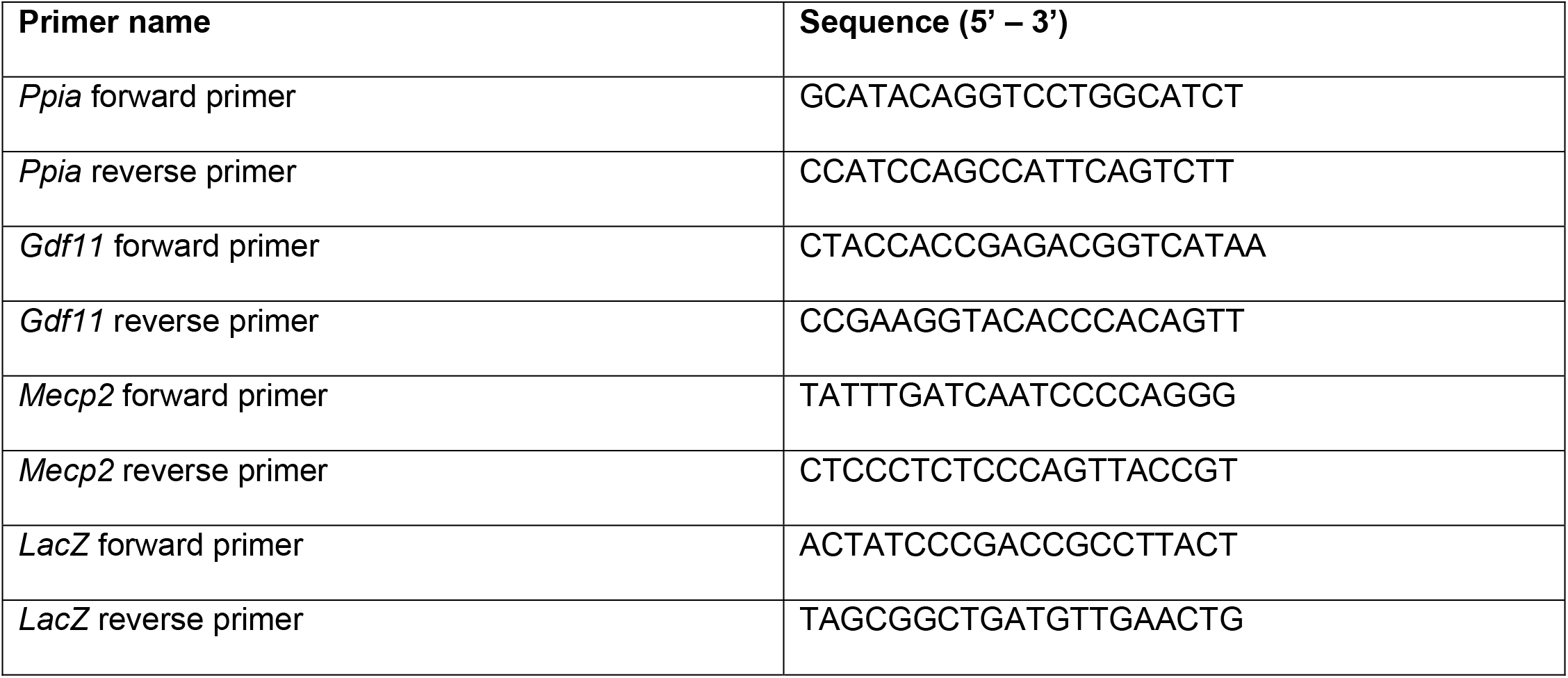

### Behavioral assays

All behavioral assays were performed during the light period. Mice were habituated to the test room for at least 30 minutes before each test. Mice were given at least one day to recover between different assays. All testing, data acquisition, and analyses were carried out by an individual blinded to the genotype.

#### Elevated Plus Maze

The lighting in the test room was set to 750 lux, and the background noise level was set to 62 dB with a white noise generator. After habituation, the mice were placed in the center zone of a plus maze that has two open arms. The mice were placed facing one of the open arms and allowed to freely move for 10 minutes. The movement of the mice was tracked with the ANY-maze software system (Stoelting Inc.). The total distance and distance traveled per zone, time spent in each arm zone, distance traveled in each arm zone, and number of zone crossings were recorded and tabulated by the software.

#### Open Field Assay

The lighting in the test room was set to 150 lux, and the background noise level was set to 62 dB with a white noise generator. After habituation, mice were placed in an open plexiglass area (40 x 40 x 30 cm), and their movement and behavior were tracked by laser photobeam breaks. Mice were allowed to freely move for 30 minutes. The total distance traveled, horizontal or vertical laser beam breaks (activity counts), entries into the center 10 x 10 cm, and time in the center 10 x 10 cm were recorded and tabulated by the AccuScan Fusion software (Omnitech Electronics Inc).

#### Light-Dark Box Assay

The lighting in the test room was set to 150 lux, and the background noise level was set to 62 dB with a white noise generator. After habituation, mice were placed in an plexiglass arena with an open zone (36 x 20 x 26 cm) and a closed, dark zone made of black plastic (15.5 x. 20 x 26 cm) with a 10.5 x. 5 cm opening. Mice were allowed to freely move for 10 minutes. The movement of the mice was tracked by laser photobeam breaks.

The total and zone-specific distance traveled, the total or zone-specific horizontal or vertical laser beam breaks (activity counts), entries into the dark zone, and time spent in either zone were recorded and tabulated by the AccuScan Fusion software (Omnitech Electronics Inc).

#### Three-chamber Assay

The lighting in the test room was set to 150 lux, and the background noise level was set to 62 dB with a white noise generator. Age and gender matched C57BL6/J mice were used as novel partner mice. Two days before the test, the novel partner mice were habituated to the wire cups (3-inch diameter by 4-inch height) for 1 hour per day for two days. After habituation, the test mice were placed in a clear plexiglass chamber (24.75 x 16.75 x 8.75 inch) with two removable partitioned that divide the chamber into three zones. Empty wire cups were placed in the left and right chambers. For the acclimation phase, the mice were placed into the center zone and the partitions removed. The mice were allowed to freely explore the apparatus for 10 minutes. For the socialization phase, a novel partner mouse was randomly placed into the left or right wire cup. An inanimate object as control was plaed in the wire cup of the opposing chamber. The mice were allowed to explore the apparatus again for 10 minutes. The total distance traveled was tracked by the ANY-maze software system (Stoelting Inc). For all phases, the total amount of time the mice spent rearing, sniffing, or pawing the cup was recorded manually.

#### Rotating rod

The lighting in the test room was at ambient levels, and the background noise level was set to 62 dB with a white noise generator. After habituation, mice were placed on an accelerating rotarod apparatus (Ugo Basile). The rod accelerated from 4 to 40 r.p.m. for 5 minutes. For cohorts containing the *MECP2-TG* allele (Figure 3), the maximum trial length was 10 min, for all other testing, the maximum trial length was 5 min. Mice were tested four trials each day, with an interval of at least 30 min between trials, for four consecutive days. The time it took for each mouse to fall from the rod (i.e., latency to fall) was recorded.

#### Fear Conditioning

Animals were habituated in an adjacent room to the testing room with a light level of 150 lux and noise level of 62 dB using a white noise generator. On the training day, mice were brought to the testing groom (ambient light, no noise) and placed into a chamber containing a grid floor that can deliver an electric shock with one mouse per chamber (Med Associates Inc). The chamber was housed within a sound-attenuating box with a digital camera, loudspeaker, and a light. The training paradigm consisted of 2 minutes of no noise or shock, then a tone for 30 seconds (5 kHz, 85 dB), ending with a foot shock for 2 seconds (1 mA). The tone and shock pairing was repeated after 2 minutes of no noise or shock. The mice were returned to their home cages after training. On the testing day, occurring 24 hours later, two tests are performed – context and cued learning. For the context test, the mice were placed in the exact same chamber with no noise or shock for 5 minutes. After one-hour post-context test, the cued learning test was performed. For the cued learning test, the mice were placed in a novel environment for 6 minutes. The first 3 minutes with no noise or shock, and the last 3 minutes with tone only (5 kHz, 85 dB). Movement was recorded on video and freezing, as defined by absence of all movement except respiration, was quantified using the FreezeFrame software (ActiMetrics) using a bout duration of 1 second and movement threshold of 10.

### Histology and staining assays

#### Cresyl violet staining

Brains from 16-week-old mice were dissected and freshly embedded and frozen in OCT. Sagittal slices were made with frozen tissues and dehydrated in 95% and 100% ethanol washes (3×2 min). Slides were washed with 1:1 chloroform:100% ethanol for 20 minutes to strip lipids. Slides were rehydrated with 95% to 70% ethanol to distilled water gradients (2 mins each). Then stain Cresyl-violet 5 mins (Solution of 0.2 % Cresyl Violet: 0.1 g Cresyl Violet; 100 ml ddH2O; 0.3 ml glacial acetic acid; 0.0205 g sodium acetate. Adjust pH 3.5). Then dehydrate slides in alcohol gradients (75%-95%-100%-100%) to xylene (2 mins each). Finally, the slides are mounted using Cytoseal 60 (VWR). The images were scanned with Zeiss Axio Scan.Z1 at 20x.

#### Brain tissue fixation and sectioning

We used design-based stereology protocol for fixing, sectioning, and quantifying proliferation markers in the subgranular zone of the dentate gyrus [Z+P citation]. Adult, 16-week-old mice (*n* =5 for *Gdf11^tm2b/+^* and *n* = 3 for wild-type littermate controls) were given a subcutaneous injection of analgesia (Buprenorphine, 0.5mg/kg) 30 minutes prior to anesthesia (Rodent Combo III, 1.5ml/kg) via intraperitoneal injection. Mice were perfused with ice-cold PBS followed by 4% PFA. Brains were removed and placed in 4% PFA overnight. Brain samples were then washed with ice-cold PBS, incubated in 15% sucrose in PBS overnight, and stored in 30% sucrose in PBS with 0.01% sodium azide. Experimenter was blind to genotype throughout the sectioning, imaging, and quantification process. Serial sections of the left hemisphere along the sagittal plane at 40um thickness were cut on a Leica SM2010 R sliding microtome and free-floating serial sections were stored in PBS + 0.01% sodium azide at 4°C.

#### EdU and Immunofluorescent staining

Mice were injected with 100 mg/kg of 5-ethynyl-2’-deoxyuridine (EdU; Invitrogen #E10187) by intraperitoneal injection for 5 consecutive days. Brains were perfused and sectioned as described above 2 days after the final injection. The Click-iT EdU Imaging Kit with AlexaFluor 647 dye (Invitrogen #C10340) was used to detect EdU in sections following manufactures protocol. Free-floating sections were stained with mouse anti-Gfap (Sigma #G3893, 1:1000 dilution) and rabbit anti-Sox2 (Abcam #ab97959, 1:500 dilution) before or after EdU Click-it reaction. Sections were washed with PBS (3X) and incubated in permeabilization buffer (0.3% Triton-X-100 in PBS) for 5 min at room temperature. Sections were placed in blocking buffer (0.3% Triton-X, 3% normal donkey serum) for 30 min at room temperature and then incubated with primary antibodies with 1% BSA in blocking buffer overnight, rocking at 4°C. Sections were washed with PBS (3X) then incubated with secondary antibodies at 1:1000 dilutions (donkey anti-rabbit Alexa 488; Jackson ImmunoResearch #711-545-152 and donkey anti-mouse Alexa 555; Thermo Scientific #A-31570) in blocking buffer with 1% BSA for 1hr at room temperature. After final washes with PBS, sections were mounted onto slides and coverslips were placed using Prolong Diamond with DAPI mounting media (Invitrogen #P36966). Slides were allowed to dry overnight. Imaging and stereology quantification Stained sections were imaged on a Leica SP8X confocal microscope. Z-stack images of the entire dentate gyrus were imaged to quantify absolute numbers of EdU^+^, Gfap^+^, and Sox2^+^ cells per section. Images were analyzed in LAS X software to measure the area and thickness of each section (t). Image J software was used to count EdU^+^, Sox2^+^, and Gfap^+^ cells (*Q*) following a standardized quantification method(Zhao and Praag, 2020). Briefly, the section sampling interval (*ssf*) was 1/6 for EdU and Gfap and 1/10 for Sox2 quantification. The area sampling fraction (*asf*)=1, and *h* =40. The following equation was used to calculate the total cell number per animal (Zhao and Praag, 2020): 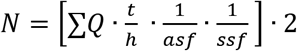.

### Statistical Analysis

Experimental analyses were performed in a blinded manner when possible. Statistical tests were performed in accordance with the experimental design. Single comparisons used Student’s t-test, whereas multi-group comparisons used one- or two-way ANOVAs as appropriate. The specific test used in each experiment is indicated in the figure legend. In each case, *, **, ***, ****, and ns denote *p* < 0.05, *p* < 0.01, *p* < 0.001, *p* < 0.0001, and *p* > 0.05, respectively.

### Resource Availability

#### Materials Availability

Mouse lines used in this study are available through Jackson Labs (*MECP2*-TG1, Jackson lab stock: 008679; *Mecp2-KO*, Jackson lab stock: 003890; or MMRRC, Stock ID: 037721).

#### Data and Code Availability

CUT&RUN raw sequencing FASTQ files are deposited to GEO (GSE #213752). This paper does not report original code. Any additional information required to reanalyze the data reported in this paper is available from the corresponding author upon request.

## ACKNOWLEDGEMENTS

We would like to thank Dr. Dah-eun Chloe Chung for critical comments of this manuscript. We further thank the Baylor College of Medicine (BCM) Genome and RNA Profiling Core and the Jan and Dan Duncan Neurological Research Institute (NRI) RNA In Situ Hybridization Core, Microscopy Core, and Animal Behavior Core. This work was supported by the Eunice Kennedy Shriver National Institute of Child Health and Human Development (NICHD) (F32HD100048 to SSB, P50HD103555 to Microscopy core, U54HD083092 to RNA In Situ Hybridization and Animal Behavior cores), Howard Hughes Medical Institute (HYZ), National Institute of Neurological Disorders and Stroke (NINDS) (R01NS057819 to HYZ, F32NS122920 to AGA), NRI Zoghbi Scholar Award through Texas Children’s Hospital (JZ). The content is solely the responsibility of the authors and does not necessarily represent the official views of the National Institutes of Health.

## AUTHOR CONTRIBUTIONS

SSB and HYZ conceived the study. SSB performed the bioinformatic analyses and experiments including molecular biology and animal work, analyzed and interpreted the data, and wrote the original manuscript. HZ supervised the study, interpreted and reviewed all the data, and edited the manuscript. AGA performed the immunofluorescence staining and quantification. JZ and MAD assisted with CUT&RUN sample generation. AJT, YWW, and ZD performed data processing of CUT&RUN raw sequencing files. All authors reviewed the manuscript and provided input.

---

**Figure S1.**
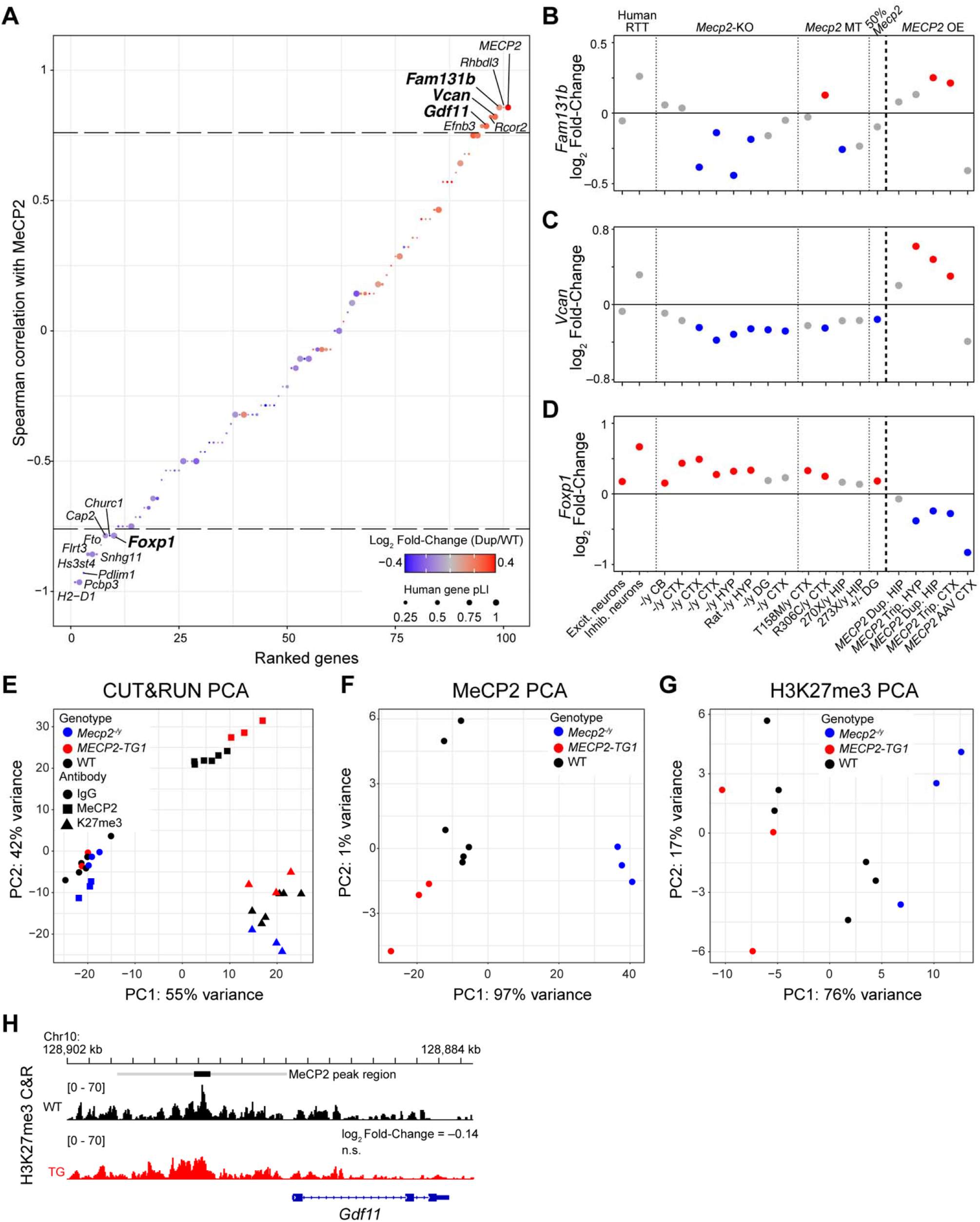
Gene expression and epigenetic changes correlated with MeCP2 protein levels. (A) Spearman correlation of genes misregulated in *MECP2* duplication mice quantifying how they track with MeCP2 levels during anti-*MECP2* antisense oligonucleotide treatment. Data points are colored by log_2_ foldchange between *MECP2* duplication and wild-type mice. The size of the dot represents the probability of loss intolerance (pLI) of the human gene if an exact match exists from the gnomad database. Genes with correlation above a threshold of 0.75 (horizontal line) are labeled by name. Genes with pLI > 0.9 are bolded. (B-C) *Fam131b, Vcan*, and *Foxp1* are MeCP2-correlated genes predicted to be intolerant to loss-of-function but are not consistently misregulated in either human Rett syndrome neurons or brain tissues of MeCP2 perturbed mouse models (see Figure 1B). Dots denote log_2_ fold-change between MeCP2 perturbed tissue and wild type, and gray color indicates *p_adj_* > 0.1. (E-I) Global assessment of MeCP2 and H3K27me3 binding in *Mecp2*-knockout, wild-type, and *MECP2-*TG1 hippocampi assessed by principal component analysis (PCA). (E) PCA of MeCP2 (square), H3K27me3 (triangle), and IgG (circle) CUT&RUN profiles in *Mecp2-*knockout (blue), wild-type (black), and *MECP2*-TG1 (red) hippocampi. (F) PCA of MeCP2 binding profiles only colored by genotype. (G) PCA of H3K27me3 binding profiles only colored by genotype. (H) CUT&RUN profiling of H3K27me3 in wild-type and *MECP2*-TG1 at the *Gdf11* locus. Black bar represents a MeCP2 peak called by MACS2 software, and the gray bar represents an expanded region of ±3.5 kb upstream of the *Gdf11* transcriptional start site. Log_2_ fold-change of binding within the gray region is shown relative to wild-type, and the *p*-value of the comparison is shown beneath the fold-change (*n* = 3 biological replicates per genotype).

**Figure S2.**
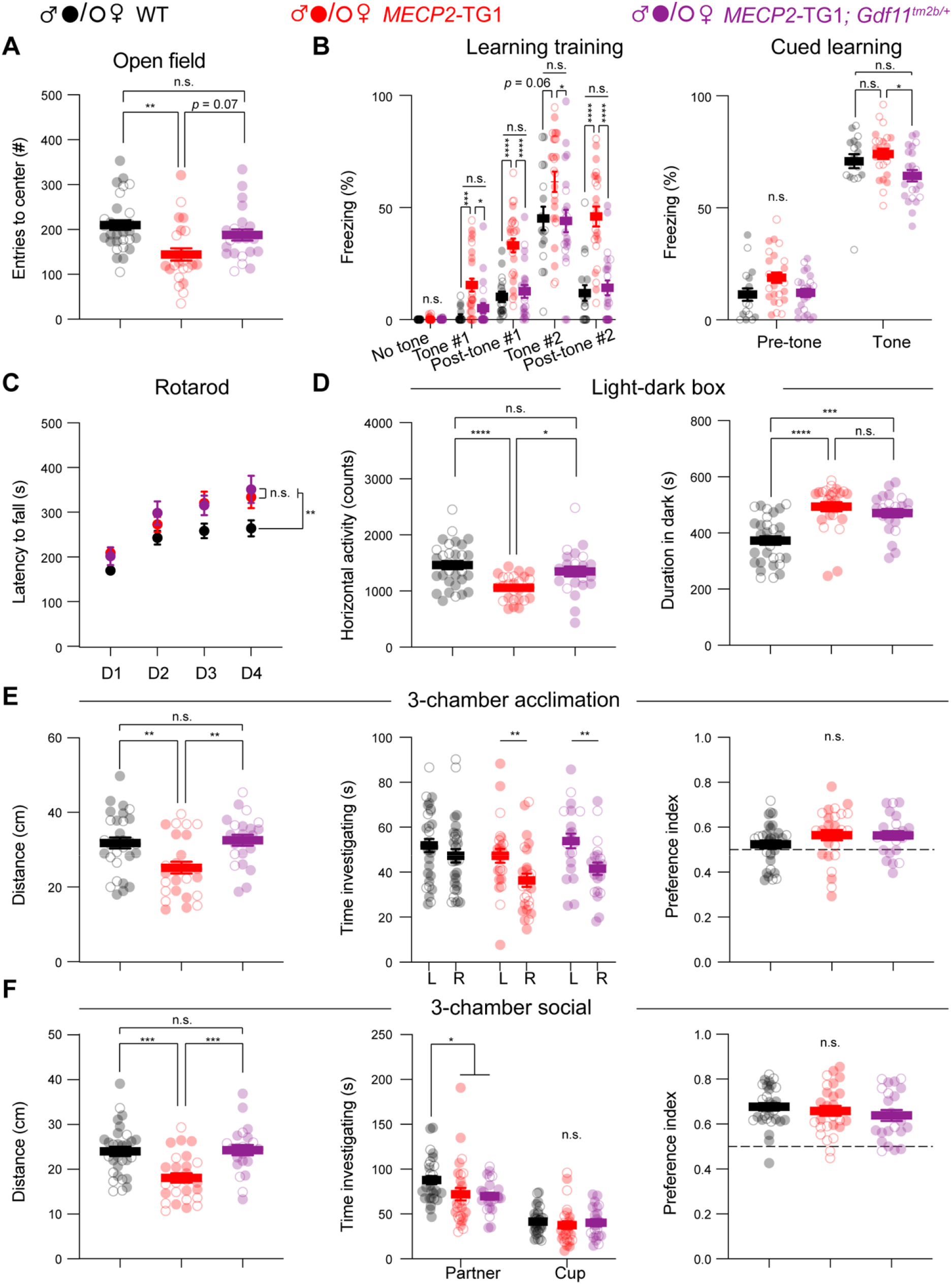
Correction of *Gdf11* dose does not ameliorate all behavioral deficits in *MECP2-TG1* mice. Behavioral characterization of *MECP2-TG1*, *MECP2-TG1; Gdf11^tm2b/+^* double mutants, and their respective wild-type littermate controls was performed beginning at 16 weeks of age. (A) Open field assessment of anxiety as measured by time in center of arena. (B) Learning assessment during training and cued fear-conditioning. Greater freezing indicates better memory of the cue. (C) Motor coordination measured by the rotating rod assay. (D) Locomotion and anxiety measured by the light-dark assay. Anxious mice prefer to spend longer duration in the dark segment of the arena. (E, F) Sociability measured by the 3-chamber assay, split into acclimation phase (E) and social phase (F). Central estimate of data is shown as mean ± sem. Closed circles denote male mouse data points and open circles denote female mouse data points. For all measurements, *n* = 30 wild-type mice (19 male, 11 female); *n* = 24 *MECP2-TG1* mice (12 male, 12 female); and *n* = 22 *MECP2-TG1; Gdf11^tm2b/+^* mice (16 male, 6 female). All data were analyzed using a Welsch one way ANOVA with Dunnett’s post hoc multiple comparisons. * *P* < 0.05, ** *P* < 0.01, *** *P* < 0.001, and **** *P* < 0.0001.

**Figure S3.**
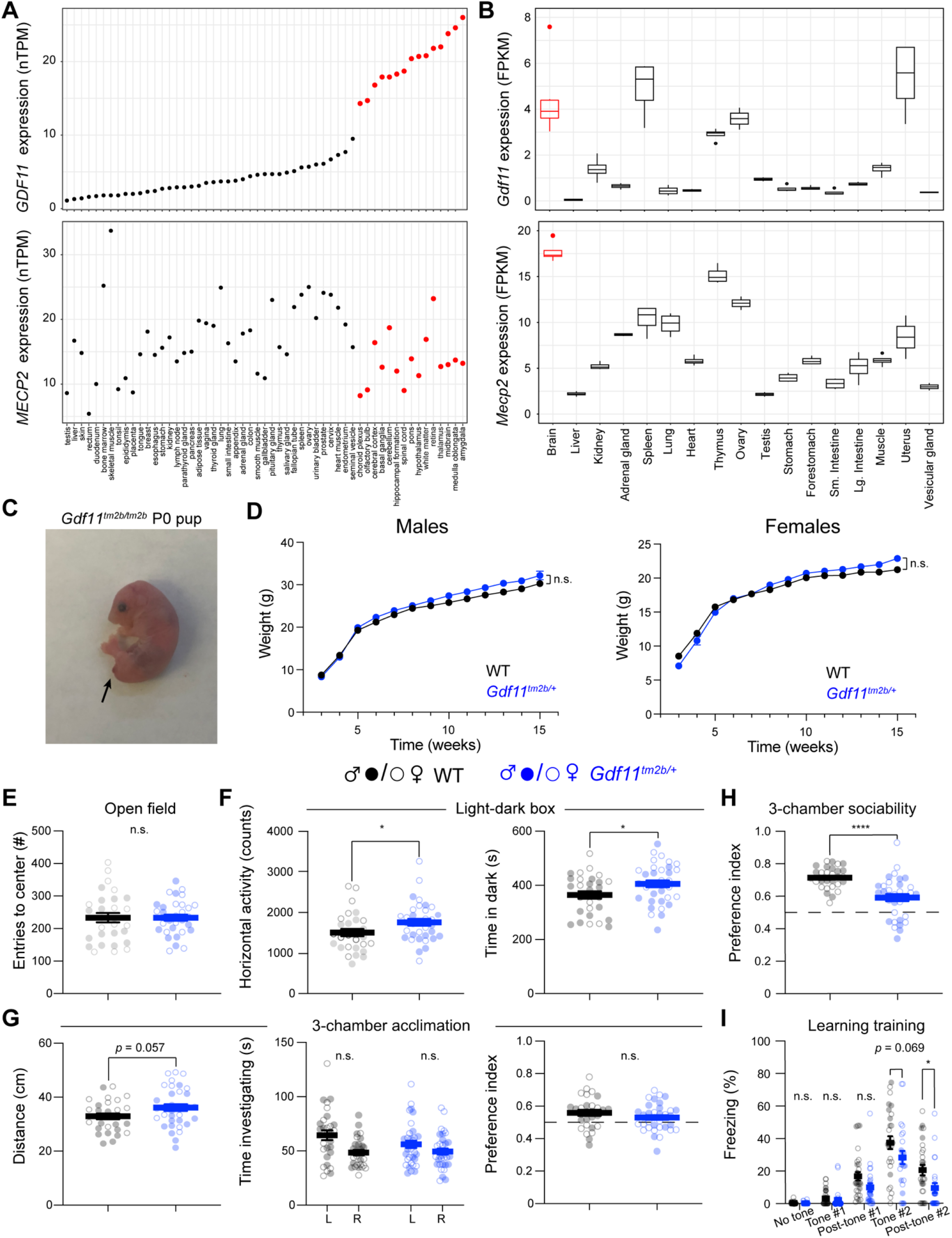
*Gdf11* is highly expressed in the brain and loss of one copy does not alter all behaviors in mice. (A) *GDF11* expression is higher the brain compared to peripheral tissues. The consensus human tissue expression dataset from the Human Protein Atlas was queried for *GDF11* (top) and *MECP2* (bottom) expression levels by tissue. Tissues are sorted based on *GDF11* expression and tissues in the central nervous system are colored red. (B) *Gdf11* is highly expressed in the mouse brain. Mouse tissue RNA-seq expression dataset was taken from Li et al. (Li et al., 2017) and queried for *Gdf11* (top) and *Mecp2* (bottom) expression levels by tissue. Brain samples are colored red. Data from *n* = 4 replicates is shown as a boxplot. (C) *Gdf11^tm2b/tm2b^* mice exhibit a truncated tail and perinatal lethality as described in the original *Gdf11^-/-^* line (McPherron et al., 1999). (D) Weight measurements in male and female *Gdf11^tm2b/+^* and wild-type littermates. Left plot displays data for males, and right plot displays data for females. (E) Open field assessment anxiety as measured by entries into the center of area. (F) Locomotion and anxiety measured by the light-dark assay. Anxious mice prefer to spend longer duration in the dark segment of the arena. (G,H) Sociability measured by the 3-chamber assay, split into acclimation phase (G) and social phase presented as the ratio of time investigating partner mouse over total time investigating (H). (I) Learning during shock-tone fear conditioning paradigm. Central estimate of behavior data is shown as mean ± sem. Closed circles denote male mouse data points and open circles denote female mouse data points. For behavioral measurements except fear conditioning, *n* = 29 wild-type mice (15 male, 14 female) and *n* = 34 *Gdf11^tm2b/+^* mice (16 male, 18 female). For fear conditioning assay (I), *n* = 28 wild-type mice (16 male, 12 female) and *n* = 24 *Gdf11^tm2b/+^* mice (9 male, 15 female). For survival analysis, *n* =15 wild-type mice and *n* = 17 *Gdf11^tm2b/+^* mice. Behavioral data were analyzed using Welsch t-test. * *P* < 0.05, ** *P* < 0.01, *** *P* < 0.001, and **** *P* < 0.0001.

**Figure S4.**
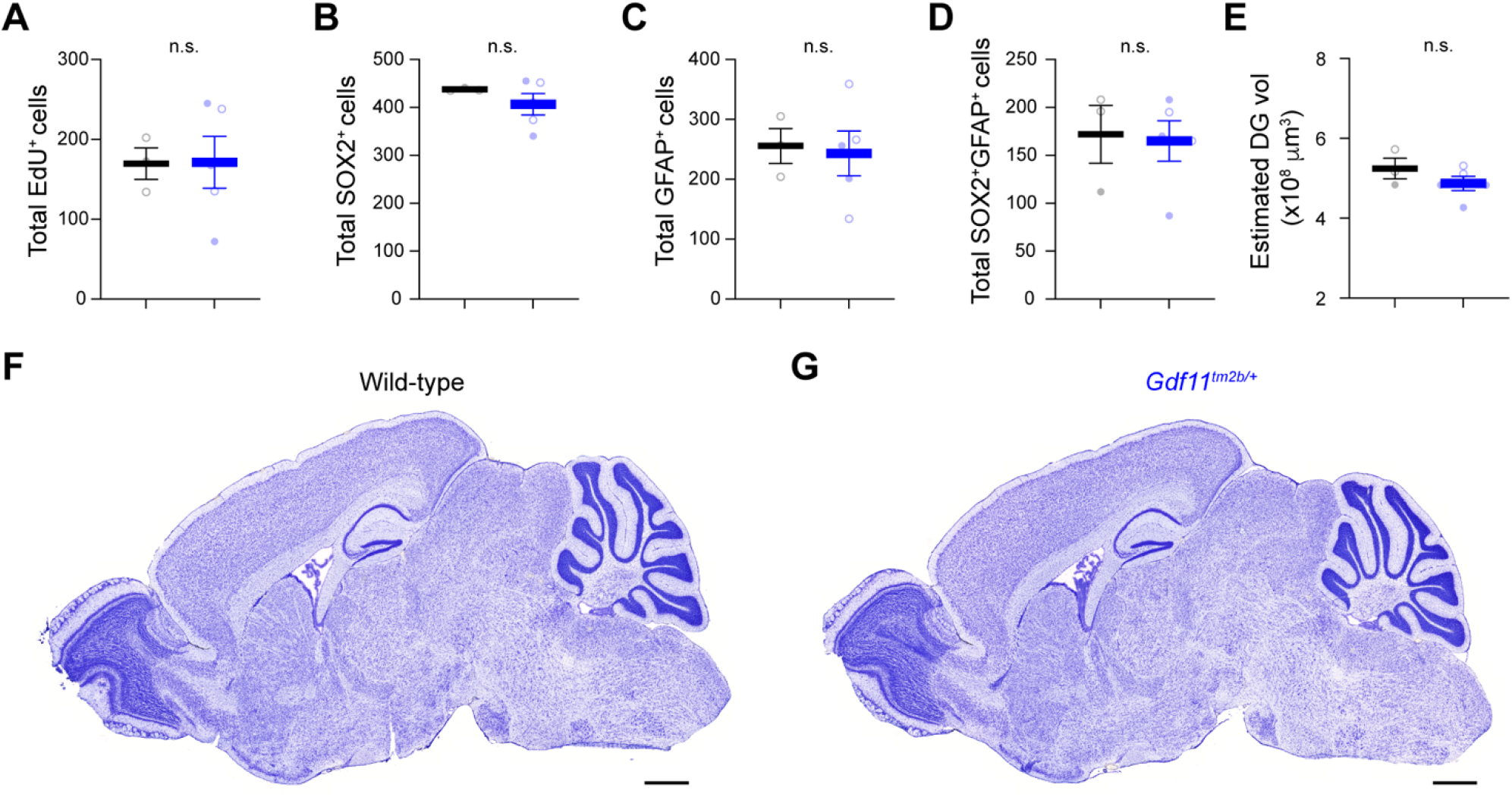
Loss of one copy of *Gdf11* does not cause gross anatomical abnormalities or gross differences in progenitor pools. (A-D) Total number of EdU (A), SOX2 (B), GFAP (C), or SOX2; GFAP double positive (D) cells scored by stereology. Counts are sums from *n* = 3-6 separate slices per animal (*N* = 3 wild-type animals and 5 *Gdf11^tm2b/+^* animals). (E) Estimated volume of the dentate gyrus from stereological quantification. (F,G) Images of Cresyl violet stain of sagittal sections from wild-type (A) or *Gdf11^tm2b/+^* brains. Scale bar is 1 mm. Each data point is the sum for one animal. Closed circles denote male animals and open circles denote female animals. Data are presented as mean ± sem. Data were analyzed using Welsch’s t-test.

**SUPPLEMENTAL TABLE S1.**
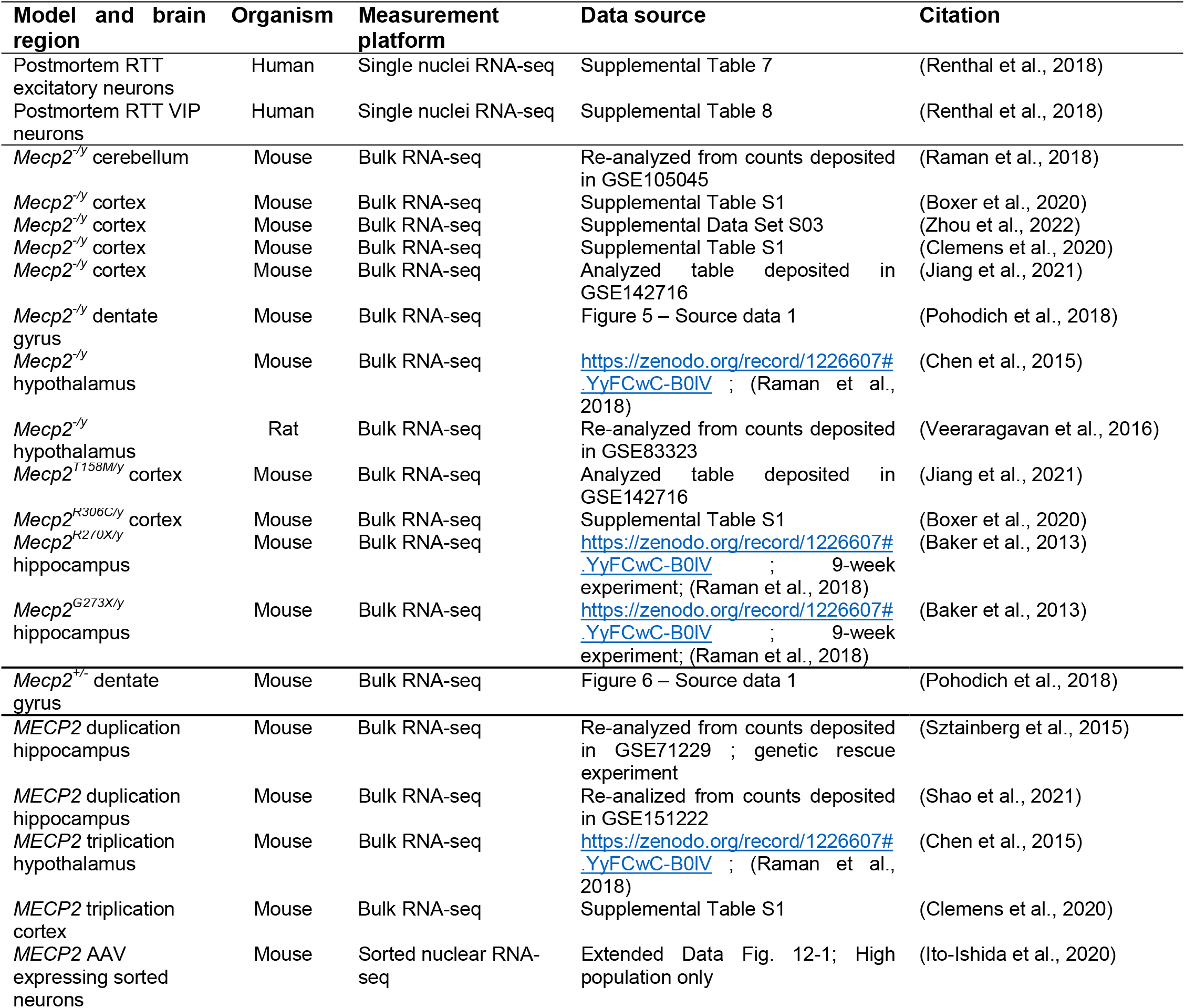

